# *In silico* characterization of *MTP1* gene associated with Zn homeostasis across different dicot plant species

**DOI:** 10.1101/2020.10.03.324863

**Authors:** Ahmad Humayan Kabir

## Abstract

Zinc (Zn) is tightly regulated in plants. The MTP1/ZAT (metal tolerance protein) plays a critical role in adjusting Zn homeostasis upon Zn fluctuation in plants. This study characterizes MTP1 homologs with particular emphasis on AtMT1 in various dicot plants. The protein BLAST search was used to identify a total of 21 MTP1 proteins. Generally, all these MTP1 proteins showed around 400 residues long, six transmembrane helices, stable instability index along with cation transmembrane transporter activity (GO:0008324). These physio-chemical features of MTP1 can be utilized as a benchmark in the prediction of Zn uptake and tolerance in plants. These MTP1 homologs were located on chromosomes 2, 7, and 14 with one exon. Motif analysis showed conserved sequences of 41-50 residues belonging to the family of cation efflux, which may be helpful for binding sites targeting and transcription factor analysis. Phylogenetic studies revealed close similarities of *AtZAT* with *Glycine max* and *Medicago trunculata* that may infer a functional relationship in Zn tolerance or uptake across different plant species. Further, interactome analysis suggests that AtZAT is closely linked cadmium/zinc-transporting ATPase and ZIP metal ion transporter, which could provide essential background for functional genomics studies in plants. The network of AtZAT is predominantly connected to cadmium/zinc-transporting ATPase (HMA2, HMA3, HMA4), cation efflux protein (MTP11), and metal tolerance protein C3 (AT4G58060). The Genevestigator platform further predicts the high expression potential of AtMTP1 in root tissue at the germination and grain filling stage. The structural analysis of MTP1 proteins suggests the conserved N-glyco motifs as well as similar hydrophobicity, net charge and nonpolar residues, alpha-helix in all MTP1 proteins. Altogether, these *in silico* characterization features of MTP1 and its orthologs will provide an essential theoretical background to perform wet-lab experiments and to better understand Zn homeostasis aiming to develop genetically engineered plants.

## 1. Introduction

Zinc (Zn) is an essential micronutrient for plants. Zn functions in photosynthetic and gene expression processes in addition to enzymatic and catalytic activities (Welch 2001). Zn deficiency resulted in a decline in stomatal activity, chlorophyll synthesis, and metabolic activity in plants (Mattiello et al. 2015; Cabot et al. 2019). The Zn is also a co-factor for transcription factors, enzymes, and protein interaction domains in Arabidopsis (Kramer, 2005). In addition, Zn ion can replace other metal ions, such as Fe, Mn, Ca, and Mg from the binding sites (Kramer, 2005; Hotz et al. 2004). In contrast, the excess accumulation of Zn ions can cause severe damage to plant cells (Dräger et al. 2004). Plants possess tightly regulated homeostasis mechanisms to maintain Zn uptake, distribution, and storage.

The AtMTP1, also known as ZAT, was the first member of the Cation Diffusion Facilitator (CDF) family members (Van der Zaal et al. 1999). Most CDF proteins have six transmembrane domains (TMDs) and a preserved C-terminal domain in the cytoplasm (Gustin et al. 2011). Among the CDF proteins, MTPs (metal tolerant proteins) are heavy metal efflux transporters in plants. *MTP* genes are not generally essential for Zn transport activity but could facilitate vacuolar sequestration of excess in the cytoplasm (Kobae et al. 2004). However, in rice, *MTP11* was found to be responsive to Zn starvation conditions (Ram et al. 2019). Most MTPs are located in the tonoplast and function as Zn and Cd antiporters involved in the sequestration or efflux of these ions to minimize metal toxicity (Kobae et al. 2004). The overexpression of *OsMTP1* in yeast and tobacco in yeast and tobacco improved Cd tolerance in rice (Das et al., 2016). Plant MTPs are grouped into seven groups, namely groups 1, 5, 6, 7, 8, 9, 12, based on annotated Arabidopsis MTP sequences (Ram et al., 2019; Migocka et al. 2015). However, the *AtMTP1* and *AtMTP3* have been shown to be associated with Zn transport in Arabidopsis. Further, both proteins function in the vacuolar sequestration of excess Zn (Desbrosses-Fonrouge et al., 2005; Arrivault et al., 2006). Studies suggest that MTP1 is Zn/H^+^ antiporter effluxing zinc out of the cytoplasm of plant cells (Kawachi et al., 2008). When ectopically overexpressed in Arabidopsis, AtMTP1 confers enhanced Zn tolerance (Van der Zaal et al. 1999).

Although *MTP1* is a crucial transporter linked to Zn homeostasis in plants, we still have limited literature on the characteristics and role of this transporter in many plant species. However, in-depth functional analysis and interactions of MTP1 with homologs remained mostly unknown. Therefore, the molecular characterization of MTP1 homologs may provide in-depth insight into these genes/proteins. In this study, we have searched Arabidopsis MTP1 (ZAT) orthologs in different plant species. The CDS, mRNA, and protein sequences of these MTP1 orthologs were analysed with advanced bioinformatics and an online-based platform.

## 2. Methods

### 2.1. Retrieval of MTP1 genes/proteins

Arabidopsis At*MTP1/ZAT* gene named as AT2G46800 in Uniprort/Aramene/Araport database (protein accession: NP_001324595.1 and gene accession: NM_001337216.1) was obtained from NCBI to use as a reference for homology search (Stephen et al. 1997). The search is limited to records that include: *Arabidopsis thaliana* (taxid:3702), *Solanum esculentum* (taxid:4081), *Brachypodium distachyon* (taxid:15368), *Oryza sativa* (taxid:4530), *Triticum aestivam* (taxid:4565), *Sorghum bicolor* (taxid:4558), *Zea mays* (taxid:4577), *Medicago truncatula* (taxid:3880), *Brassica oleracea* (taxid:3712), *Glycine max* (taxid:3847), *Beta vulgaris* (taxid:161934), *Pisum sativum* (taxid:3888), *Nicotiana tabacum* (taxid:4097), *Solanum tuberosum* (taxid:4113), *Setaria italica* (taxid:4555), in which results were filtered to match records with expect value between 0 and 0.

### 2.2. Analyses of MTP1 genes/proteins

Physico-chemical features of MTP protein sequences were analyzed by the ProtParam tool (https://web.expasy.org/protparam) as previously instructed (Gasteiger et al. 2005). Chromosomal and exon position was detected by the ARAMEMNON database (http://aramemnon.uni-koeln.de/). The CELLO (http://cello.life.nctu.edu.tw) server predicted the subcellular localization of proteins (Yu et al. 2006). Protein domain families were searched in the Pfam database (http://pfam.xfam.org), and functions were assessed by the Phytozome v12.1 database (El-Gebali et al. 2019. The structural organization of MTP1 genes was predicted by FGENESH online tool (Solovyev et al. 2016).

### 2.3. Phylogenetic Relationships and Identification of Conserved Protein Motifs

Multiple sequence alignments of MTP1 proteins were performed to identify conserved residues by using Clustal Omega. Furthermore, the five conserved protein motifs of the proteins were characterized by MEME Suite 5.1.1 (http://meme-suite.org/tools/meme) with default parameters, but five maximum numbers of motifs to find (Timothy et al. 1994). Motifs were further scanned by MyHits (https://myhits.sib.swiss/cgi-bin/motif_scan) web tool to identify the matches with different domains (Sigrist et al. 2010). The MEGA (V. 6.0) developed the phylogenetic tree with the maximum likelihood (ML) method for 1000 bootstraps using 21 MTP1 homologs from 17 plant species (Tamura et al. 2013).

### 2.4. Interactions and co-expression of MTP1 protein

The interactome network of AtMTP1 protein was generated using the STRING server (http://string-db.org) visualized in Cytoscape (Szklarczyk et al. 2019). Further, gene network, co-occurrence, and neighborhood pattern were also retried from the STRING server. Additionally, the expression data of Arabidopsis MTP1 was retrieved from Genevestigator software and analyzed at hierarchical clustering and co-expression levels based on the Affymetrix genome array.

### 2.5. Structural analysis of MTP1 proteins

Structural analysis, such as transmembrane domains and Helicoidal representation, was constructed with Protter (http://wlab.ethz.ch/protter/start) tool (Omasits et al. 2014) and HeliQuest (https://heliquest.ipmc.cnrs.fr/) server (Gautier et al. 2008). Lastly, a two-dimensional secondary structure of MTP1 proteins constructed GORIV (https://npsa-prabi.ibcp.fr/NPSA/npsa_gor4.html).

## 3. Results

### 3.1. Retrieval of MTP1 transporter genes/proteins

Arabidopsis *AtMTP1*, as referred to as *ZAT*, was searched against 15 species in the NCBI database to get the FASTA sequence of the protein (NP_001324595.1) and mRNA (NM_001337216.1). This particular gene/protein is also named as AT2G46800 in Uniprort/Aramene/Araport database. The blast analysis of MTP1 protein showed 21 orthologs of the cation efflux family by filtering the E-value to 0.0. The retrieved proteins include 2 proteins for *A. thaliana*, 3 proteins for *Brassica oleracea*, 2 proteins for *Solanum tuberosum*, 1 protein for *Solanum lycopersicum*, 5 proteins for *Glycine max*, 4 proteins for *Nicotiana tabacum*, and 4 proteins for *Medicago truncatula* (Table 1).

**Table 1.** List of MTP1 homologs and their physio-chemical features.

| No. | NCBI entry | Species | Molecular Function | Gene/protein features of retrieved sequences |  |  |  |  |  |  |
| --- | --- | --- | --- | --- | --- | --- | --- | --- | --- | --- |
|  |  |  |  | Localization | Protein length | MW (Da) | pI | transmembrane helices (TMH) | Instability index | Grand average of hydropathicity (GRAVY) |
| 1 | Protein:<br>NP_001318436.1<br>Gene:<br>NM_001337216.1 | *Arabidopsis thaliana* | cation transmembrane transporter activity (GO:0008324) | Vacuole | 398 | 43827.34 | 6.13 | 6 | 32.63 (stable) | 0.147 |
| 2 | Protein:<br>AAD11757.1<br>Gene:<br>AF072858.1 | *Arabidopsis thaliana* | cation transmembrane transporter activity (GO:0008324) | Vacuole | 398 | 44063.70 | 6.05 | 6 | 34.46 (stable) | 0.166 |
| 3 | Protein:<br>XP_013636161.1<br>Gene:<br>XM_013780707.1 | *Brassica oleracea* var. oleracea | cation transmembrane transporter activity (GO:0008324) | Vacuole | 381 | 42175.56 | 5.68 | 6 | 36.23 (stable) | 0.192 |
| 4 | Protein:<br>VDD16513.1<br>Gene:<br>LR031873.1 | *Brassica oleracea* | cation transmembrane transporter activity (GO:0008324)<br>protein binding (GO:0005515) | Vacuole | 861 | 94711.75 | 6.14 | 6 | 43.63 (unstable) | -0.242 |
| 5 | Protein:<br>XP_013637363.1<br>Gene:<br>XM_013781909.1 | *Brassica oleracea* var. oleracea | cation transmembrane transporter activity (GO:0008324) | Vacuole | 381 | 41965.44 | 5.89 | 6 | 38.32 (stable) | 0.235 |
| 6 | Protein:<br>XP_006359546.1<br>Gene:<br>XM_006359484.2 | *Solanum tuberosum* | cation transmembrane transporter activity (GO:0008324) | Vacuole | 385 | 42861.52 | 5.86 | 6 | 31.06 (stable) | 0.162 |
| 7 | Protein:<br>XP_004242700.1<br>Gene:<br>XM_004242652.3 | *Solanum lycopersicum* | cation transmembrane transporter activity (GO:0008324) | Vacuole | 416 | 46022.69 | 6.00 | 6 | 27.65 (stable) | 0.007 |
| 8 | Protein:<br>NP_001242638.2<br>Gene: | *Glycine max* | cation transmembrane transporter activity (GO:0008324) | Vacuole | 408 | 45248.03 | 6.23 | 6 | 33.30 (stable) | 0.059 |
| 9 | Protein:<br>NP_001312370.1<br>Gene: | *Nicotiana tabacum* | cation transmembrane transporter activity (GO:0008324) | Vacuole | 418 | 46210.91 | 6.00 | 6 | 29.86 (stable) | -0.002 |
| 10 | Protein:<br>ACU18393.1 | *Glycine max* | cation transmembrane transporter activity (GO:0008324) | Vacuole | 397 | 43887.62 | 6.26 | 6 | 33.12 (stable) | 0.130 |
| 11 | Protein:<br>XP_016493500.1 | *Nicotiana tabacum* | cation transmembrane transporter activity (GO:0008324) | Vacuole | 418 | 46105.81 | 6.02 | 6 | 28.23 (stable) | 0.022 |
| 12 | Protein:<br>ACU21236.1 | *Glycine max* | cation transmembrane transporter activity (GO:0008324) | Vacuole | 419 | 46118.79 | 6.12 | 6 | 31.26 (stable) | 0.051 |
| 13 | Protein:<br>BAD89563.1 | *Nicotiana*<br>*tabacum* | cation<br>transmembrane<br>transporter<br>activity<br>(GO:0008324) | Vacuole | 418 | 46204.94 | 6.07 | 6 | 28.89<br>(stable) | 0.012 |
| 14 | Protein:<br>NP_001242882.2 | *Glycine max* | cation<br>transmembrane<br>transporter<br>activity<br>(GO:0008324) | Vacuole | 419 | 46176.83 | 6.08 | 6 | 31.50<br>(stable) | 0.043 |
| 15 | Protein:<br>XP_014623915.1 | *Glycine max* | cation<br>transmembrane<br>transporter<br>activity<br>(GO:0008324) | Vacuole | 422 | 46558.32 | 6.10 | 6 | 31.35<br>(stable) | 0.049 |
| 16 | Protein:<br>NP_001312828.1 | *Nicotiana*<br>*tabacum* | cation<br>transmembrane<br>transporter<br>activity<br>(GO:0008324) | Vacuole | 418 | 46142.83 | 6.04 | 6 | 28.66<br>(stable) | 0.010 |
| 17 | Protein:<br>RHN73387.1 | *Medicago*<br>*truncatula* | cation<br>transmembrane<br>transporter<br>activity<br>(GO:0008324) | Vacuole | 408 | 45185.71 | 5.89 | 6 | 31.79<br>(stable) | 0.057 |
| 18 | Protein:<br>XP_013444540.1 | *Medicago*<br>*truncatula* | cation<br>transmembrane<br>transporter<br>activity<br>(GO:0008324) | Vacuole | 385 | 42458.02 | 6.02 | 6 | 31.15<br>(stable) | 0.201 |
| 19 | Protein:<br>XP_024627417.1 | *Medicago*<br>*truncatula* | cation<br>transmembrane<br>transporter<br>activity<br>(GO:0008324) | Vacuole | 386 | 42589.22 | 6.02 | 6 | 31.02<br>(stable) | 0.205 |
| 20 | Protein:<br>XP_003594932.1 | *Medicago*<br>*truncatula* | cation<br>transmembrane<br>transporter<br>activity<br>(GO:0008324) | Vacuole | 407 | 45054.52 | 5.89 | 6 | 31.92<br>(stable) | 0.053 |
| 21 | Protein:<br>XP_006359535.1 | *Solanum*<br>*tuberosum* | cation<br>transmembrane<br>transporter<br>activity<br>(GO:0008324) | Vacuole | 415 | 45972.68 | 6.05 | 6 | 28.74<br>(stable) | 0.005 |

### 3.2. Physiochemical features and localization of MTP proteins

In total, 21 MTP1 homologs were found by homology quest in proteome datasets of 15 plant species. They encoded a protein with residues of 398–419 amino acids having 41965.44 to 46558.32 (Da) molecular weight, and 5.68 to 6.26 pI value, 27.65 to 43.63 instability index, and −0.002 to 0.235 grand average of hydropathicity (Table 1). Notably, all these MTP1 proteins showed 6 transmembrane helices (TMH). The subcellular localization of MTP1 homologs was predicted as the vacuole. In addition, all these homologs show cation transmembrane transporter activity as a molecular function (Table 1). ARAMEMNON analysis showed that MTP1 homologs were located at chromosomes 2, 7, and 14 in which exon was located at 1828-3024, 1911-3041, and 1818-2999 base pair, respectively (Table 2). In addition, the structural analysis of the MTP1s gene showed the presence of 1 exon in homologs (Table 2). The position of transcriptional start site (TSS) ranged from 53-330, whereas the coding sequences were located as early as 13 to 2223 base positions. The PolA is consistently positioned after the coding region in all MTP1 genes showing the position at 1434-2274 (Table 2).

**Table 2.** Organization of MTP1 genes and position features.

| No. | Gene Accession | Chromosome<br>number | Number of exon and<br>position | Position of<br>Transcriptional Start<br>Site (TSS) | Position of<br>Coding sequence | Position of<br>PolA |
| --- | --- | --- | --- | --- | --- | --- |
| 1 | NM_001337216.1 | 2 | 1 (1828-3024) | 330 | 1027 - 2223 | 2274 |
| 2 | AF072858.1 | 2 | 1 (1828-3024) | 53 | 214 - 1410 | 1461 |
| 3 | XM_013780707.1 | 2 | 1 (1828-3024) | 130 | 291 - 1436 | 1494 |
| 4 | LR031873.1 | 2 | 1 (1828-3024) | - | 78 - 389 | - |
| 5 | XM_013781909.1 | 2 | 1 (1828-3024) | 151 | 224 - 1369 | 1434 |
| 6 | XM_006359484.2 | 7 | 1 (1911-3041) | - | 113 - 1270 | - |
| 7 | XM_004242652.3 | 7 | 1 (1911-3041) | 167 | 387 - 1637 | 1710 |
| 8 | NM_001255709.3 | 14 | 1 (1818-2999) | - | 215 - 1441 | 1474 |
| 9 | NM_001325441.1 | 7 | 1 (1911-3041) | - | 1 - 1257 | - |
| 10 | NM_001255709.3 | 14 | 1 (1818-2999) | - | 215 - 1441 | 1474 |
| 11 | NM_001325899.1 | 7 | 1 (1911-3041) | 100 | 253 - 1458 | 1711 |
| 12 | NM_001255953.3 | 14 | 1 (1818-2999) | - | 148 - 1407 | 1599 |
| 13 | NM_001325899.1 | 7 | 1 (1911-3041) | 100 | 253 - 1458 | 1711 |
| 14 | NM_001255953.3 | 14 | 1 (1818-2999) | - | 148 - 1407 | 1599 |
| 15 | NM_001255953.3 | 14 | 1 (1818-2999) | - | 148 - 1407 | 1599 |
| 16 | NM_001325899.1 | 7 | 1 (1911-3041) | 100 | 253 - 1458 | 1711 |
| 17 | CM010649.1 | 14 | 1 (1818-2999) | - | 13 - 225 | - |
| 18 | XM_024771650.1 | 14 | 1 (1818-2999) | - | 159 - 1316 | - |
| 19 | XM_024771650.1 | 14 | 1 (1818-2999) | - | 159 - 1316 | - |
| 20 | XM_024776611.1 | 14 | 1 (1818-2999) | - | 368 - 1591 | 1678 |
| 21 | XM_006359474.2 | 7 | 1 (1911-3041) | - | 311 - 1558 | 1801 |

### 3.3. Conserved motif, Sequence similarities, and phylogenetic analysis

We have used the MEME tool to search for the five most conserved motifs in identified 21 MTP1 homologs (Table 3). Motifs 1, 2, 3, and 5 were 50 long residues of amino acids, while motif 4 was 41 long amino acids. All motifs relating to the family of MTP1 proteins are present in all MTP1 sequences. The analysis showed that motif 1 (DAAHLLSDVAAFAISLFSLWAAGWEATPRQSYGFFRIEILGALVSIQMIW), 2 (WYKPEWKIVDLICTLIFSVIVLGTTINMJRNILEVLMESTPREIDATKLE), 3 (HIWAITVGKVLLACHVKIRPEADADMVLDKVIDYIKREYNISHVTIQIER), 4 (DAZERSASMRKLCIAVVLCVIFMTVEVVGGIKAN), and 5 (LAGILVYEAIARLIAGTGEVDGFLMFLVAAFGLVVNJIMALLLGHDHGH) were shown the best match to cation efflux family (Table 3). Long preserved residues may also indicate highly conserved MTP transporter structures among various species. All five motifs were shared by all 21 MPT proteins among the 15 plant species (Fig. 1).

**Table 3.**
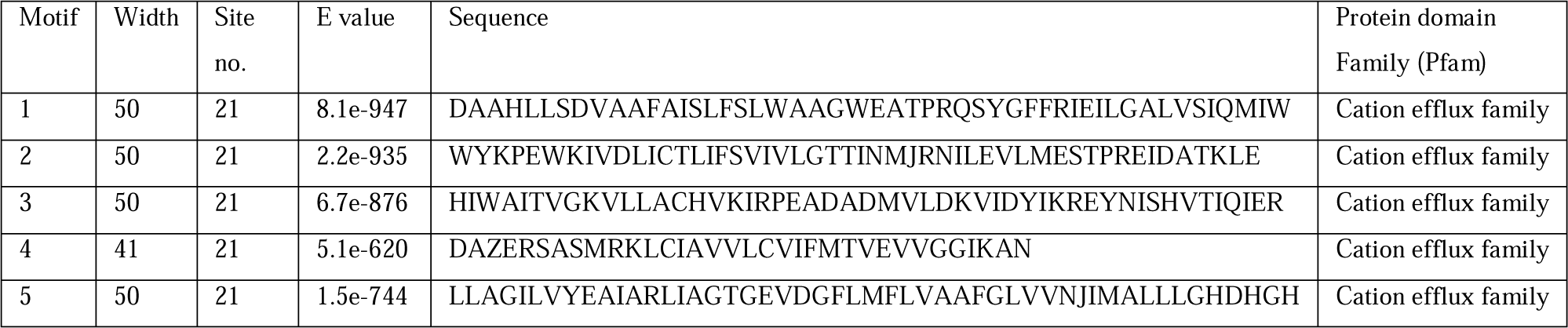
Most conserved five motifs of MTP1 homologs in 15 plant species.

**Fig. 1.**
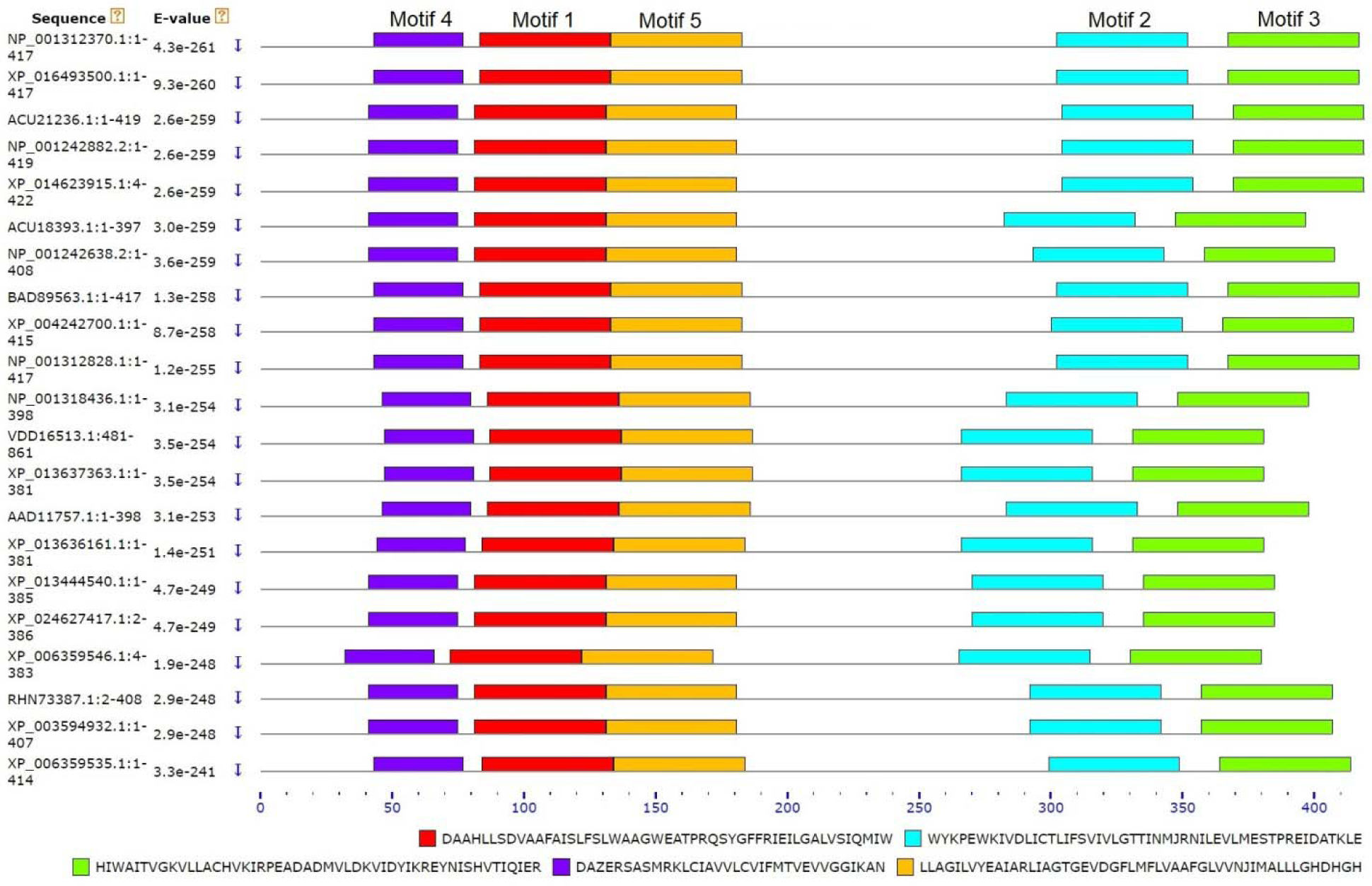
Location of five motif in 21 MTP proteins of 15 plant species.

To classify other preserved protein regions, we aligned all 21 MTP1 transporter sequences by Clustal Omega (Supplementary Fig. S1.). The MTP1 proteins showed 70% to 100% similarities among the different plant species. The consensus sequence ranged from 70%-100% (Supplementary Fig. S1.). The phylogenetic was divided into two main groups based on tree topologies, such as A, B, C, D, E, F, and G (Fig. 2). In group A, 4 MTP proteins of *Nicotiana tabacum* and 1 MTP1 of *Solanum lycopersicum* have formed a cluster. Group B consisted of 2 MTP1 proteins of *Brassica oleracea*, 1 of *Solanum tuberosum*, and 1 of *Glycine max*. Two MTP proteins of *Arabidopsis thaliana* and *Medicago trunculata* formed groups C and D, respectively (Fig. 2). In group E, 4 MTP1 proteins of *Glycine max* and 2 of *Medicago trunculata* formed the cluster. Group F and G include a predicted MTP1 protein of *Solanum tuberosum* and an unnamed protein sequence of *Brassica oleracea*, respectively (Fig. 2). In this phylogenetic tree, two MTP1 proteins of *Arabidopsis thaliana* and two predicted MTP1 proteins of *Brassica oleracea* showed the highest bootstrap value (99%).

**Fig. 2.**
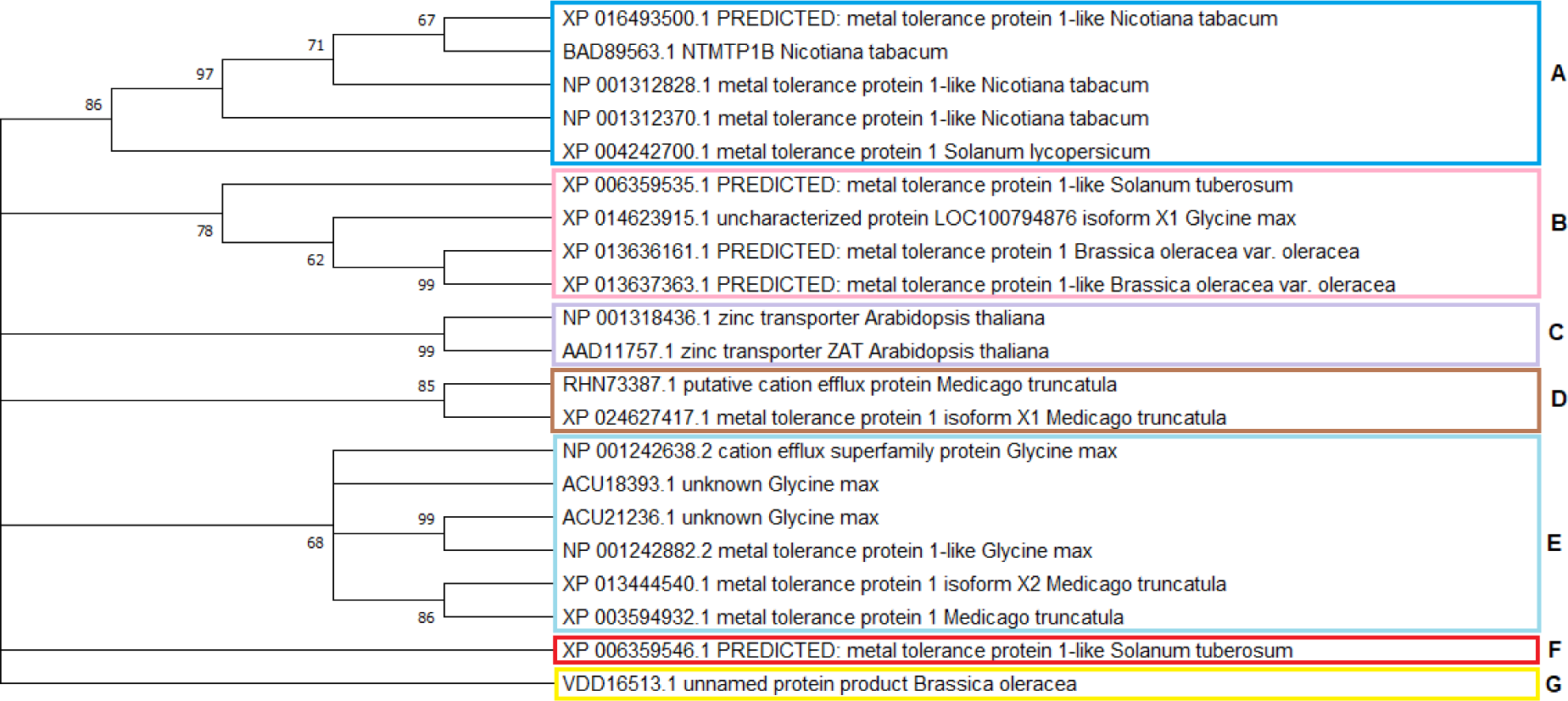
Phylogenetic tree of 21 MTP1 homolog proteins using Mega 6. Statistical method: Maximum likelihood phylogeny test, test of phylogeny: bootstrap method, No. of bootstrap replications: 1000.

### 3.5. Predicted interaction partner analysis

Predicted interaction partner analysis was performed for AtMTP1/AtZAT (AT2G46800). STRING showed ten putative interaction partners of a zinc transporter (ZAT) and cation diffusion facilitator (CDF), which include HMA2, HMA3, HMA4, IAR1, ZIP9, NRAMP3, RNR1, MTP11, AT1G51610, and AT3G58060 (Fig. 3). Among them, HMA2, HMA3, and HMA4 are responsible for cadmium/zinc ATPase. MTP11 and AT1G51610 are attached to the cation efflux family. Further, RNR1, IAR1, NRAMP3, ZIP9, and AT3G58060 are linked to ribonucleoside-diphosphate reductase large subunit, IAA-alanine resistance protein 1, natural resistance-associated macrophage protein 3, ZIP metal ion transporter family, and putative metal tolerance protein C3, respectively (Fig. 3a). Gene network analysis showed a close association of *AtZAT* with some genes associated with metal transport and tolerance, which includes *HMA2* (cadmium/zinc-transporting ATPase HMA2), *HMA3* (putative inactive cadmium/zinc-transporting ATPase HMA3), *HMA4* (putative cadmium.zinc-transporting ATPase HMA4), *MTP11* (cation efflux family protein involved in Mn tolerance) and AT4G58060 (putative metal tolerance protein C3 involved in metal sequestration) genes (Fig. 3b). Further, MTP1/ZAT protein and its partners of *Arabidopsis thaliana* showed close co-occurrence with *Arabidopsis lyrata, Capsella, Camelina sativa*, and *Brassica species* (Fig. 4). These species are also the close neighborhoods of Interaction Partner proteins (Fig. 4).

**Fig. 3.**
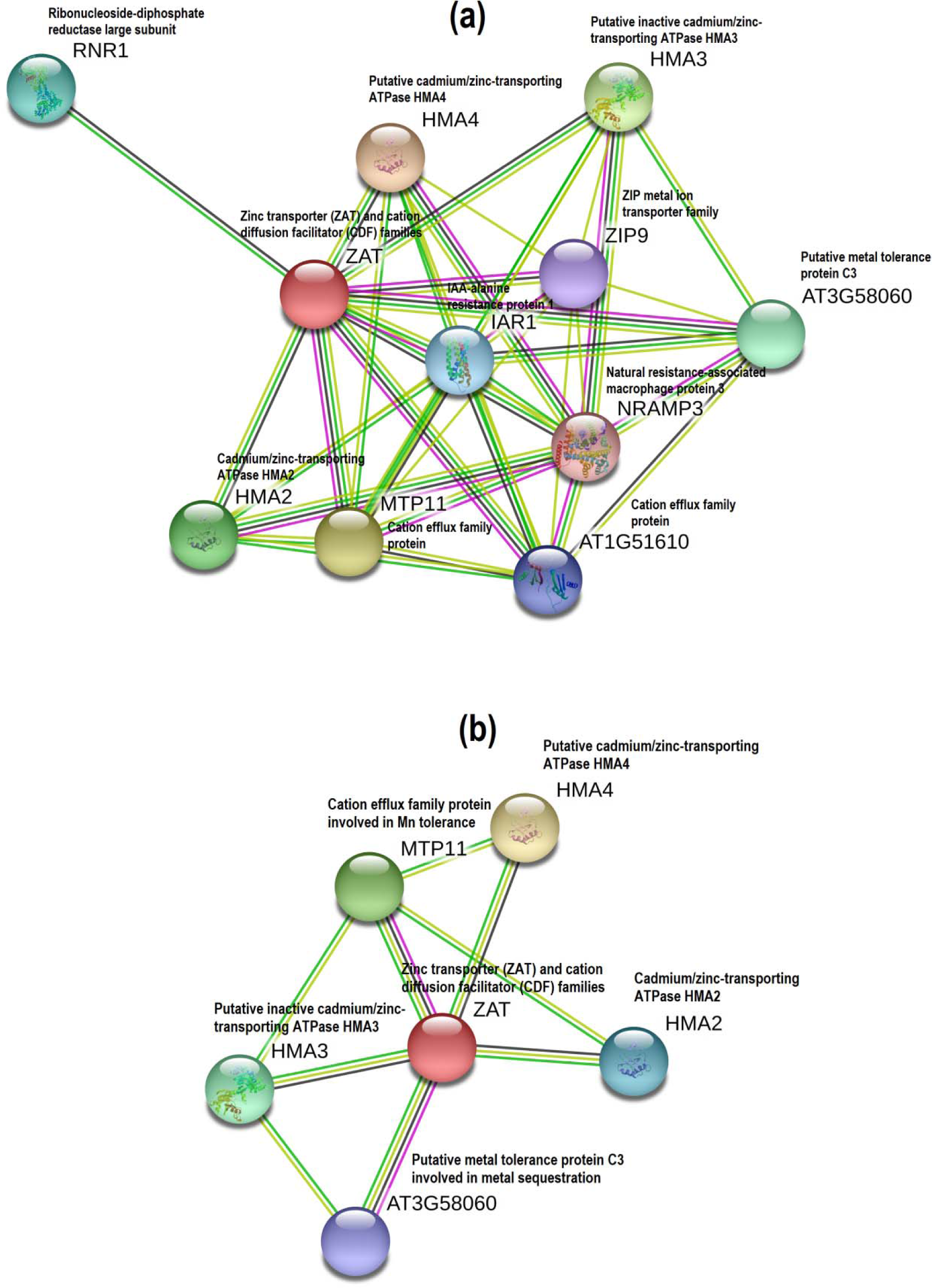
Predicted gene interaction partners (a) and (b) networks of AtMTP1/AtZAT protein. Interactome was generated using Cytoscape for STRING data.

**Fig. 4.**
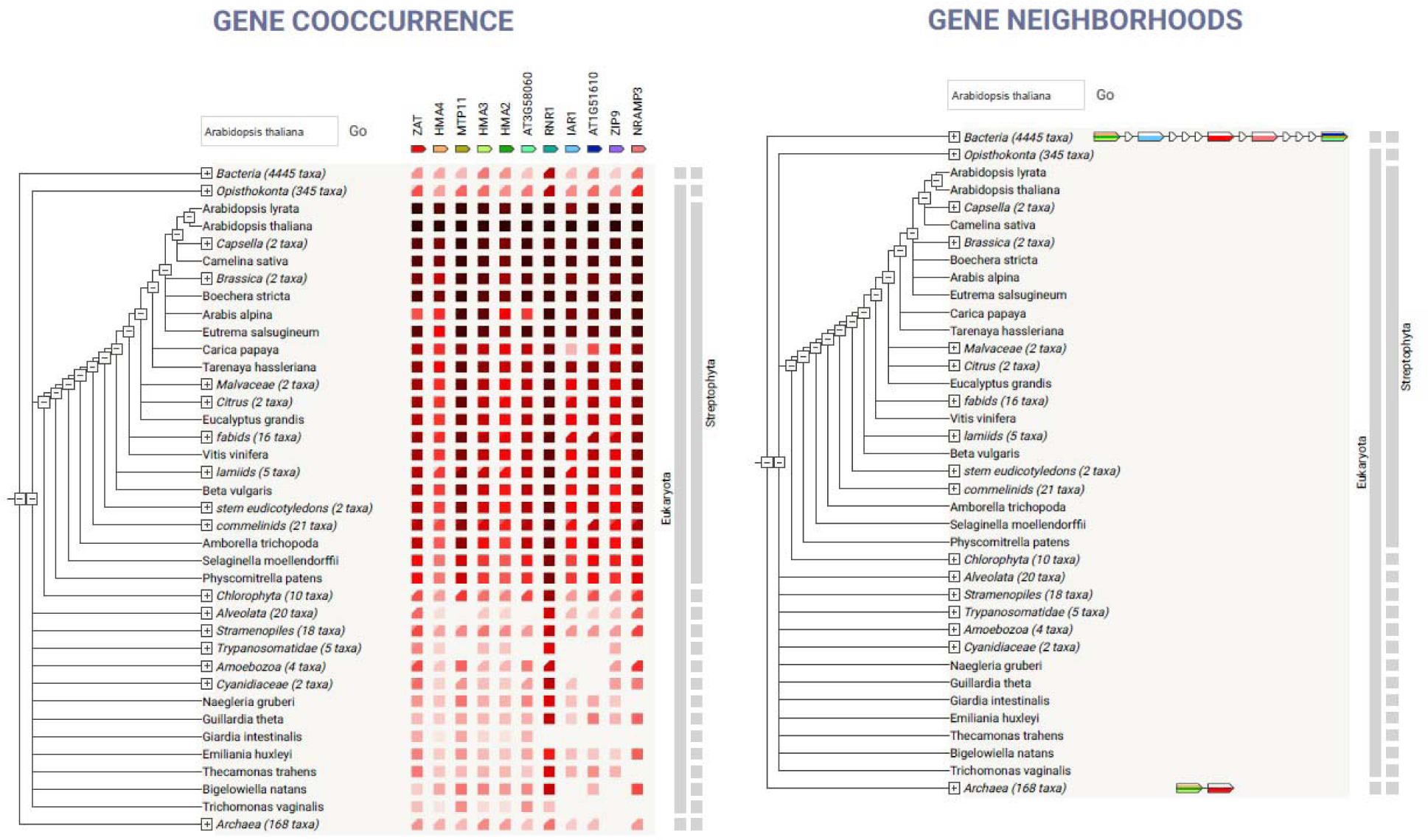
Co-occurrence and neighborhoods of Predicted interaction partners of AtMTP1/AtZAT protein.

The genvestigator analysis against Affymetrix Arabidopsis ATH1 genome array showed co-expression data of *MTP1* in different anatomical parts, perturbations, and developmental stages (Fig. 5). In the anatomical part, the *MTP1* was found to be highly co-expressed in the apical root meristem. Subsequently, *MTP1* showed strong co-expression in root cortex protoplast, root epidermis and quiescent center protoplast, root epidermis, and lateral root cap protoplast and root tip (Fig. 5a). Genes co-expressed under perturbation correlating above 0.415 showed 11 matches, which include *SKIP1, PDS1, ABF1, SBP1, SKP2A, AT3G04350, BAM1, MAX2, NUDT15, TLP1*, and *AT1G21780* (Fig. 5b). Also, MTP1 was found to be highly co-expressed in most of the developmental stages, of which germination and grain stage are two top matches were found (Fig. 5c).

**Fig. 5.**
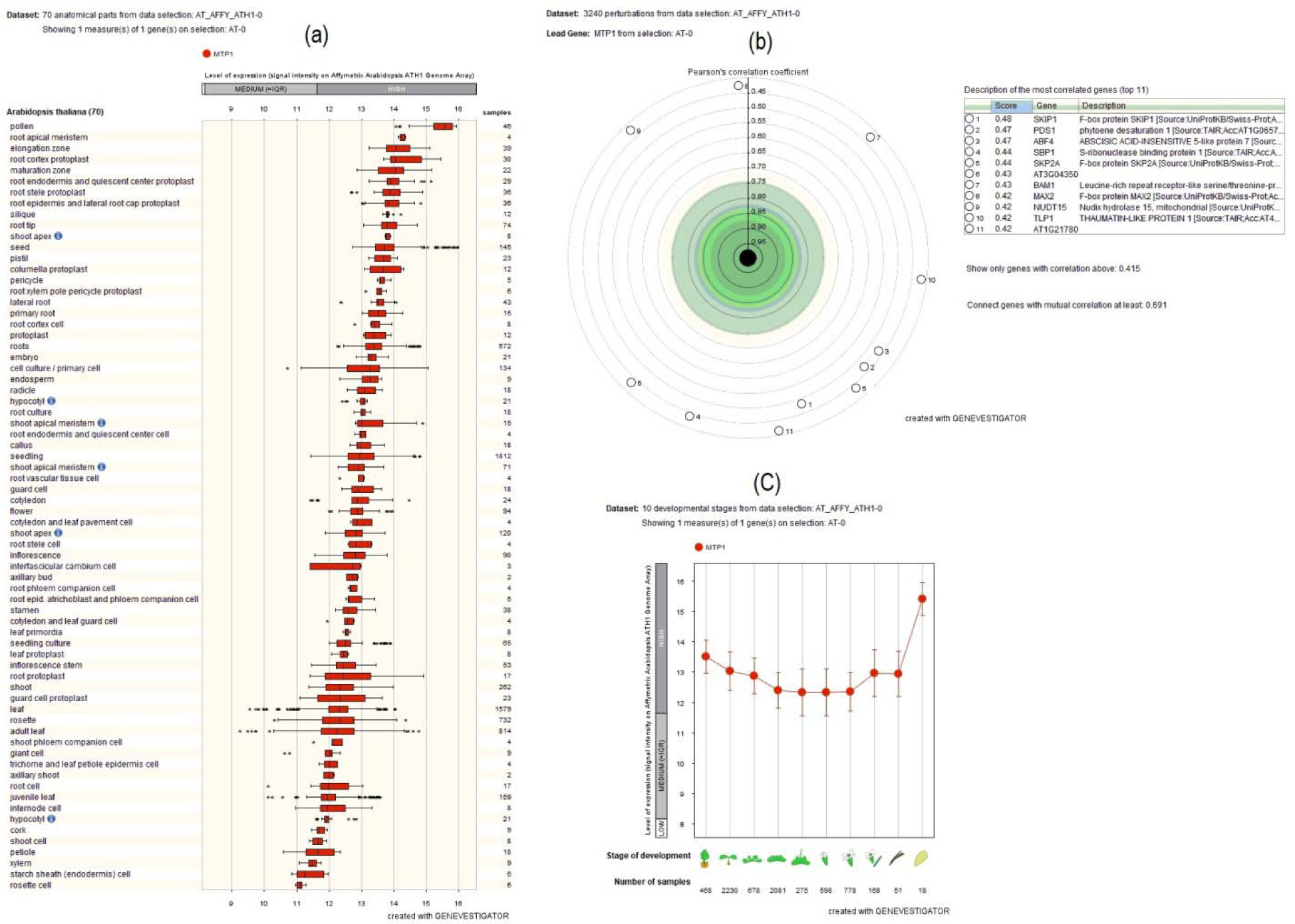
Co-expression of MTP1 in different anatomical part, perturbations and developmental stage.

### 3.6. Analysis Secondary Structure of MTP1 proteins in different plant species

Topological prediction analyses of transmembrane (TM) domains of MTP1s showed 1-6 TM domains in protein representative from each of the plant species (Fig 6). The MTP1 TM domains are well preserved in different sequences; however, amino acid sequences vary at N-termini (Fig. 6). Helical wheel representation displayed no significant variations other than the position of C and N terminus in these MTP1 proteins. Further, polar and nonpolar residues ranged from 11-12 and 6-7. All these MTP1 proteins contained special residues CYS and PRO (Supplementary Fig. S2). In addition, secondary structure prediction showed that all MTP1 proteins have above 35% α-helices, above 35% random coils and around 20% extended strands, and 40% random coils (Supplementary Fig. S3).

**Fig. 6.**
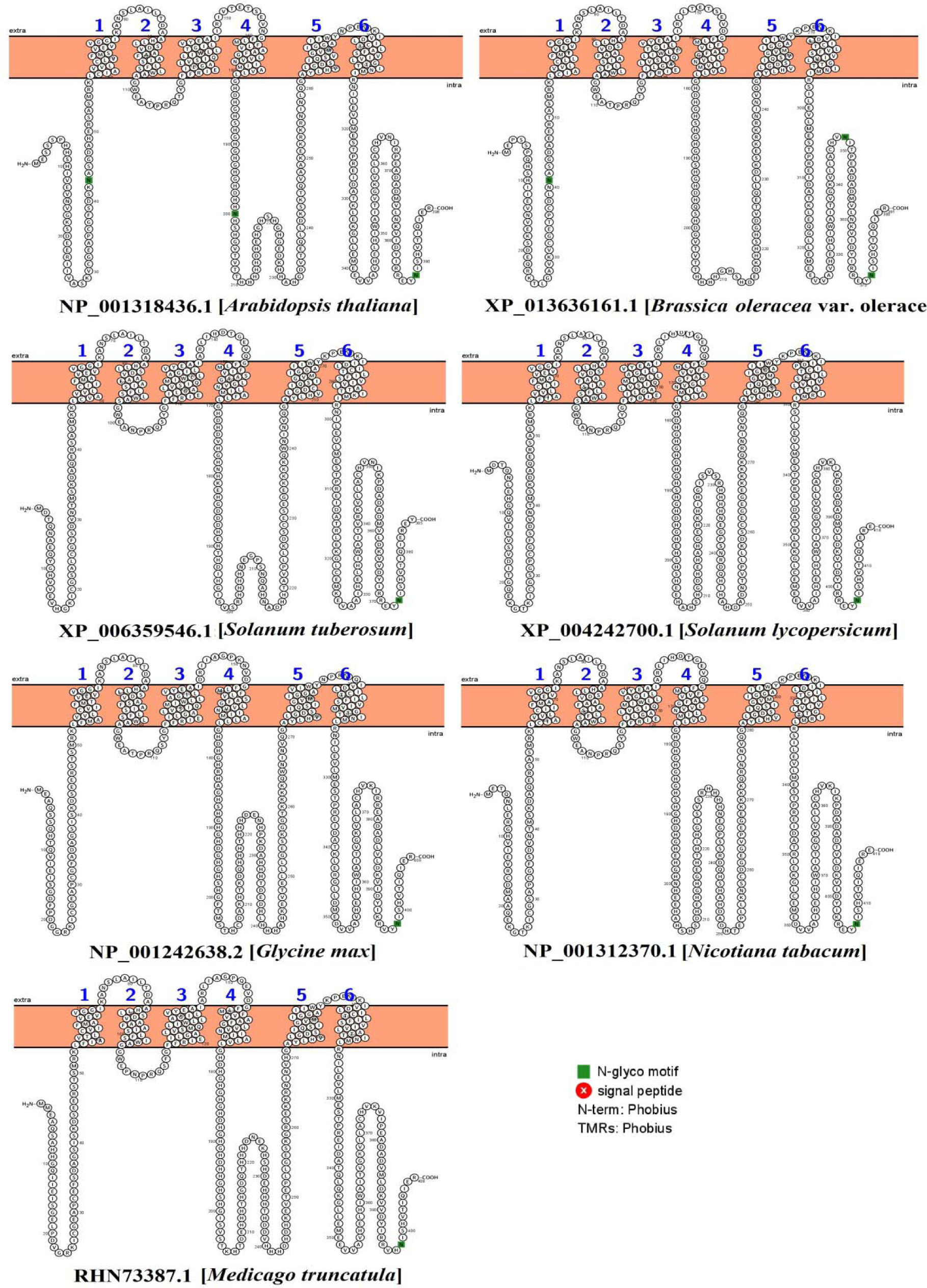
Structural analysis of MTP1 proteins in different plant species in constructed with Protter.

## 4. Discussion

Characterization of a gene is of great interest to further accelerate the wet-lab experiments in plant science. This *in silico* work, led to the identification of 21 MTP proteins among seven plant species. MSA shows these proteins are 98.5-100% coverage of similarity with 70-100% matching of consensus sequences among the different MTP proteins. Most CDF proteins contain six transmembrane domains (Wei and Fu, 2005), and all members have a characteristic C-terminal efflux domain (Maser et al., 2001). Our analysis also showed the existence of 6 TMD in all 21 MTP1 proteins connected to the cation transmembrane transporter activity (GO: 0008324).

The position or organization of the coding sequence of a gene is considered to be a critical factor in predicting evolutionary relations among the orthologues and paralogues. In this study, all MTP1 genes among the seven plant species showed 1 exon, suggesting that these MTP1 genes are phylogenetically closer to each other. Our analysis further explored the position of TSS and PolA of several MTP1 proteins, which are crucial to understanding transcriptional and post-transcriptional modification of mRNA. In general, it is known that genes without intron have recently evolved (Deshmukh et al. 2015). The subcellular localization of these MTP1 proteins was predicted as vacuoles. MTP proteins are vacuolar transporters and can isolate metals in cells (Gustin et al. 2011). However, *AtMTP1* has been shown to have Zn transport activity as well (Desbrosses-Fonrouge et al., 2005; Bloss et al., 2002). Similarly, *AtMTP3* can transport Zn and Co when expressed in the yeast mutant (Arrivault et al., 2006). Interestingly, Peiter et al. (2007) demonstrated that *AtMTP11* localized neither to vacuole or plasma membrane, but to a Golgi compartment providing tolerance to Mn. In this study, all of the identified sequences of MTP1 demonstrated acidic character having the pI value of around 6 along with the positive hydropathicity except two sequences. The protein length of 20 MTP1 proteins was 381-419, while only the MTP1 of *Brassica oleracea* (LR031873.1) showed 861 amino acids. Several studies reported the length of amino acid residues in the ZIP family transporter from 309–476 (Guerinot, 2000).

Conserved motifs are identical sequences across species that are maintained by natural selection. A highly conserved sequence is of functional roles in plants and can be a useful start point to start research on a particular topic of interest (Wong et al. 2015). Among the 21 MTP1 proteins, we searched for five motifs using the MEME tool. All of these five motifs belonged to the cation efflux family. In our search, motif 1, 2, 3, and 5 displayed 50 amino acid residues long, while motif 4 showed 41 residues long. The presence of common and long conserved residues pinpoints that MTP1 homologs may possess highly conserved structures between species. Additionally, this information can be targeted for sequence-specific binding sites and transcription factor analysis.

In phylogenetic analysis, we clustered the tree in 7 sub-groups. According to the tree, two Arabidopsis MTP1 proteins clustered within group C as expected. These AtMTP1 proteins showed the closest phylogenetic relationship with *Glycine max*, and *Medicago trunculata* MTP1 proteins resulted in 99 bootstraps. It also appears that AtMTP1 is relatively distantly related to *Nicotiana tabacum, Solanumber tuberosum, Solanum lycopersicum*, and *Brassica oleracea*. The relationship with cluster C and F was further figured out in MSA similarities index. Consistently, AtMTP1 proteins (NP_001318436.1 and AAD11757.1) demonstrated 93-97.1% similarities to the MTP1 proteins of *Glycine max* and *Medicago trunculata*. The AtMTP1 has shown to be involved with Zn tolerance (Kobae et al. 2004) and Zn transport in Arabipssis (Arrivault et al., 2006). However, it is not yet reported whether MTP1 is also involved in Zn homeostasis in other closely related plant species. Thus, our results might infer a functional relationship MTP1 sequences in Zn or other metals tolerance or uptake across different plant species.

Interactome map and neighborhood analysis were performed using the *AtAMTP1* (ZAT) (AT2G46800/NP_001324595.1/NM_001337216.1). In the interactome map, cation efflux family protein MTP11, putative cadmium/zinc-transporting ATPase HMA4, cadmium/zinc-transporting ATPase HMA2, IAA-alaline resistant protein IAR1, and ZIP metal ion transporter ZIP9 were predicted among the interaction partners of AtAMTP1. Studies demonstrated that *MTP11* plays a critical role in Mn homeostasis in rice (Zhang and Liu, 2017) and Arabidopsis (Delhaize et al., 2007). In plants, Zn homeostasis is closed associated with P-type ATPase heavy metal transporters (HMA). Both HMA2 and HMA4 were reported to be involved with Zn homeostasis in Arabidopsis (Hussain et al., 2004). ZIP family members have also been characterized in plants involved in metal uptake and transport, including Zn (Kavitha et al., 2015). Auxin participates in many plant developmental processes and stress tolerance in plants. Interestingly, the IAR1 gene, responsible for auxin metabolism, has detectable sequence similarity to a family of metal transporters (Lasswell et al., 2000). Network analysis reveals the association of cadmium/zinc transporter, cation efflux protein, and metal tolerance protein C3 with AtMTP1. We further searched for the co-occurrence and neighborhoods of AtMTP1. These analyses displayed that most nearby co-occurrence and neighborhood of the AtMTP1 gene are HMA4, MTP11, HMA3, HMA3, AT3G58060, RNR1, IAR1, AT1G51610, ZIP9, and NRAMP3 genes of *Arabidopsis lyrata* and *Calsella* sp. Overall, this interactome findings might provide essential background for functional genomics and hormone studies in plants.

The potentiality of expression of a gene in different conditions is a crucial factor in the genome editing program. The *in silico* analysis of expression profile using Affymetrix Genome Array in Genevestigator online platform showed impressive results concerning different anatomical, perturbations, and developmental stages. In this analysis, the *AtMTP1* is predominantly expressed in the different parts of root tissue, by which plants acquire metals from the soil. Several CDF and ATPase family transporters were shown root-specific expression regulating Zn and Cu homeostasis in plants (Seigneurin-Berny et al. 2005; Desbrosses-Fonrouge et al. 2005). Given the involvement of root organelle, this study further advances our knowledge to elucidate the uptake and mobilization of Zn and other metals in plants. Also, environmental stimuli or perturbations do have a strong influence on gene expression patterns in plants. Our perturbations analysis showed several correlated genes of *AtMTP1*, including *SKIP*1, *PDS1, ABF4, SBP1*, SKP2A, etc. Again, seedling and grain maturation stages were found to be highly dominant in expressing the AtMTP1 gene in Arabidopsis. These messages may provide an outline in functional genomics studies in Arabidopsis or closely related species in metal studies. Among the MTP1 protein family studied in this study showed three N-glyco motifs in *Arabidopsis thaliana* and *Brassica oleracea*, while the rest of the species showed only one. However, MTP1 revealed 6 TM located within the helices of all MTP1 proteins. In the helicoidal structure, all these MTP1 proteins showed similar hydrophobicity, net charge, and nonpolar residues. Two-dimensional structures of these MTP1 proteins consistently showed similar alpha helix, extended strand, and random coil. These findings may provide insights into the protein architecture and particular function.

## Conclusion

In conclusion, this bioinformatics analysis analyzed 21 MTP1 protein homologs in different plant species. The study showed similar physicochemical properties, gene organization, and conserved motifs related to the cation efflux family. Sequence homology and phylogenetic tree showed the closest evolutionary relationship of Arabidopsis MTP1 with *Glycine max* and *Medicago trunculata*. In addition, the interactome map displayed the co-expression of AtMTP1 with a number of closely related genes involved in Cd/Zn transport in plants. It was also predicted that *AtMTP1* is highly expressed in root tissue at early germination or grain maturation stages. Similar protein architecture and the structural organization further suggest the unique feature of this MTP1 protein across the dicot plant species. These findings will provide basic theoretical knowledge for future studies on the understanding of gene function and protein features of genes related to Zn homeostasis in various plants

---

**Supplementary Fig. S1.**
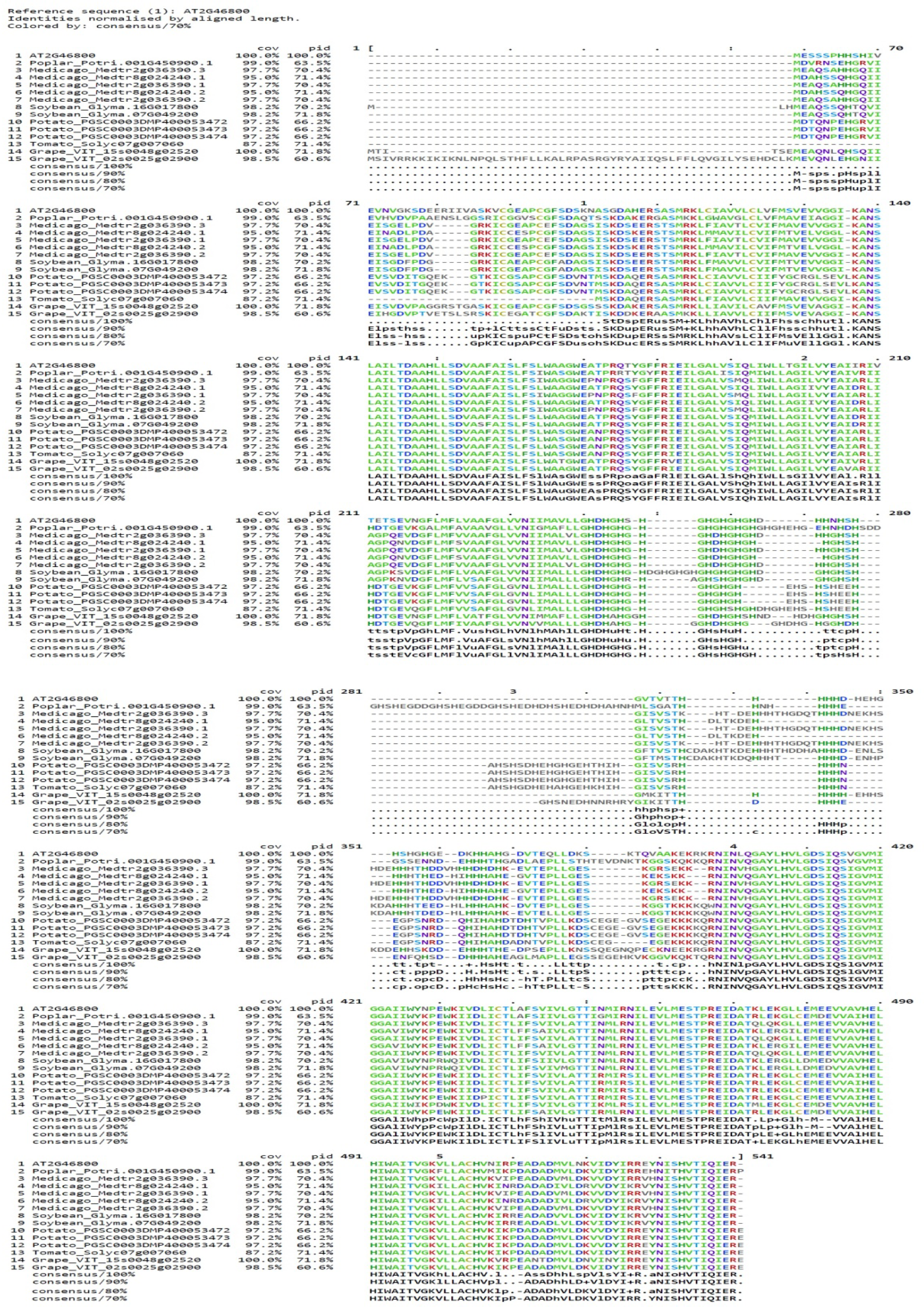
Multiple sequence alignment (MSA) of MTP1 across plant species.

**Supplementary Fig. S2.**
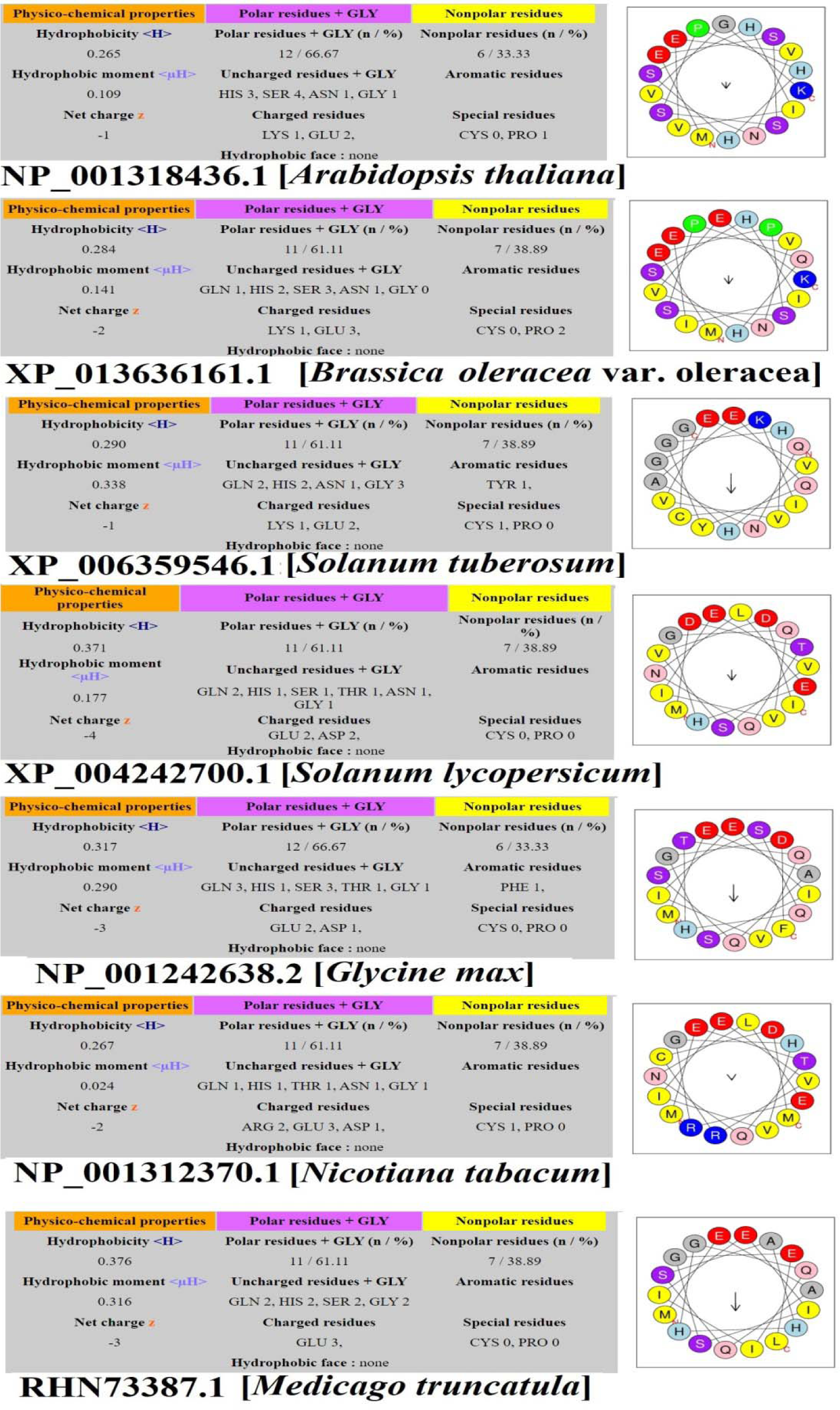
Helicoidal representation of MTP1 proteins constructed with Heliquest.

**Supplementary Fig. S3.**
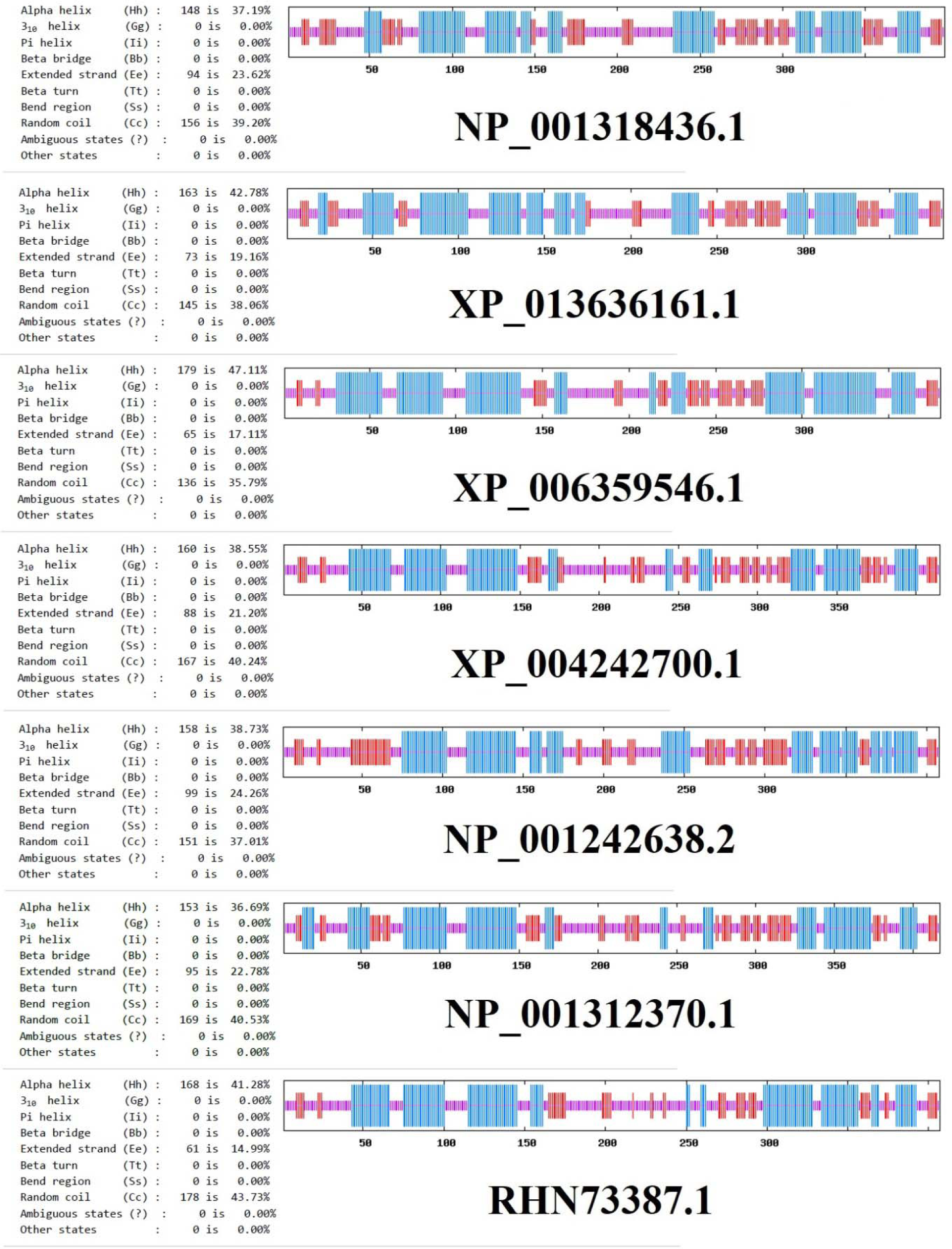
Two dimensional secondary structure of MTP1 proteins in different plant species in constructed with GORIV.

